# Transplanting enzyme active site geometry into antibody CDRs for catalytic antibody design

**DOI:** 10.64898/2026.07.25.740676

**Authors:** Yaojun Zhu

## Abstract

Antibodies provide programmable molecular recognition, whereas enzymes enable repeated chemical transformation. Catalytic antibodies seek to combine these properties within a single protein scaffold. However, conventional approaches based on transition state analogue immunisation, library screening or local mutagenesis provide limited control over the atomic arrangement of catalytic residues. They also frequently produce antibodies that bind substrates without supporting efficient chemical turnover. Recent advances in generative protein design have enabled the construction of antibody complementarity determining regions and the scaffolding of functional motifs under structural constraints. A systematic strategy for transferring experimentally supported enzyme active site geometry into antibody variable domains is still lacking. Here, we present a computational framework that treats antibody and enzyme structures as distinct but complementary inputs. Developable Fv or VHH structures provide the immunoglobulin scaffold. Enzyme complexes containing substrates, products or transition state analogues provide catalytic residues, ligand conformations, metals, cofactors and key water networks. The selected catalytic atoms are mapped into antibody complementarity determining regions, while the surrounding loops are reconstructed using antibody compatible representations and constrained all atom diffusion. Sequence design and structural back prediction are followed by filters for antibody folding, catalytic geometry, ligand positioning, conformational stability and developability. The framework avoids direct fusion of intact enzymes and antibodies. Instead, it transfers only the local geometry required for catalysis. This separation of scaffold selection from catalytic motif selection creates a testable route for determining whether natural enzyme chemistry can be embedded within antibody formats. It also provides a practical basis for evaluating substrate binding, chemical conversion, product release and catalytic turnover as separate design objectives.

**Graphical Abstract:** 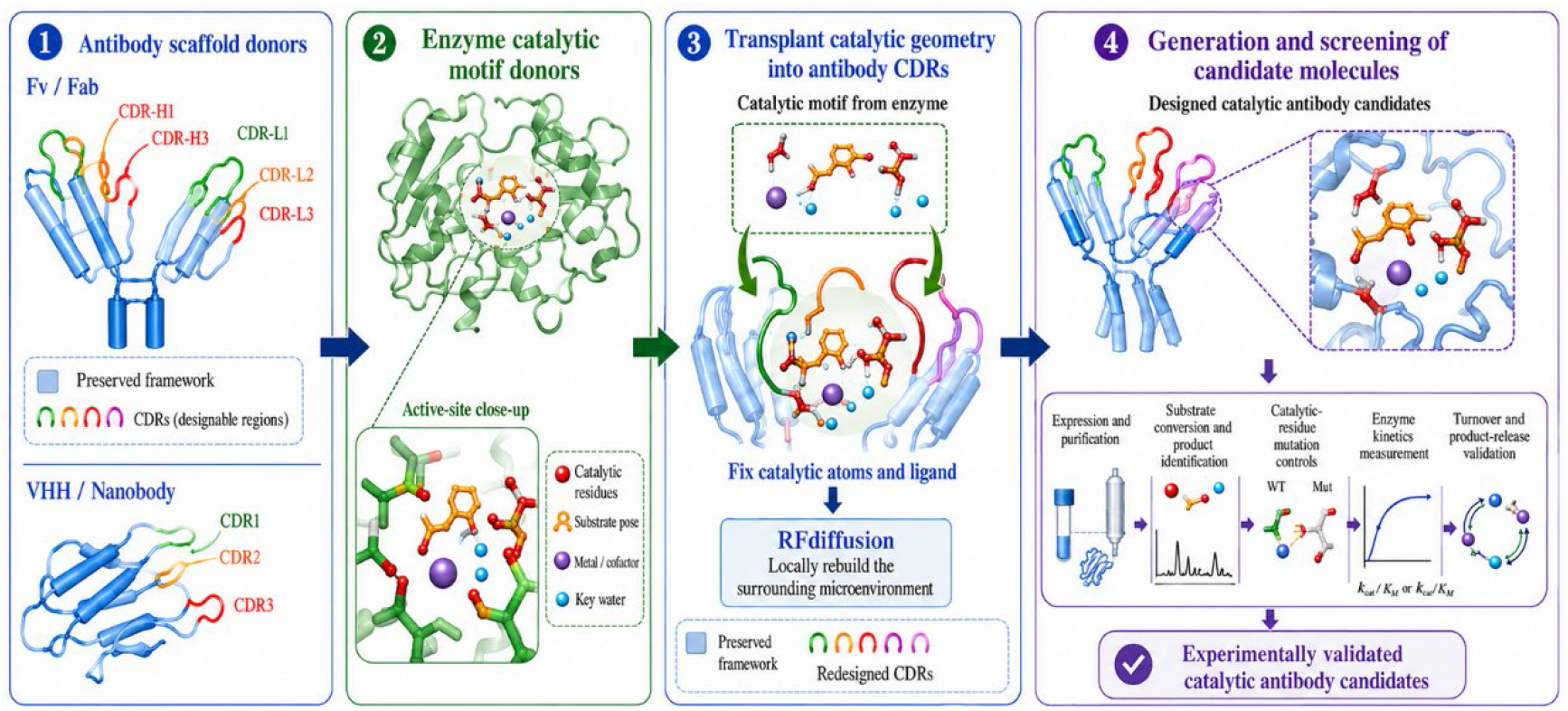

## Introduction

The feasibility of antibody-mediated catalysis was demonstrated in the 1980s. Early studies generated monoclonal antibodies by immunising animals with stable analogues of reaction transition states and found that some antibodies could accelerate specific chemical reactions [1, 2]. However, conventional catalytic antibody development first requires the artificial synthesis of specific transition-state analogues as haptens for animal immunisation [3]. The subsequent screening process is time-consuming, labour-intensive and has a low success rate [1]. Moreover, the catalytic activity of these antibodies is generally far lower than that of natural enzymes [4]. Difficult product release can also cause occupancy-based inhibition, which markedly slows the overall reaction cycle [5]. Antibodies are also large molecules with complex structures, making extensive modification difficult [6]. The commercial production cost of catalytic antibodies offers no clear advantage [7]. Together, these limitations have prevented the field from developing a mature design and application platform.

Researchers have subsequently developed antibody–enzyme fusion proteins and other conjugated systems to combine the targeting ability of antibodies with the catalytic function of enzymes [8]. However, such modular assemblies usually increase molecular weight and structural complexity. They may also introduce problems such as linker instability, difficult expression and mutual interference between functional activities [9].

In recent years, generative protein design has provided new possibilities for addressing these limitations [10]. In this study, we propose an RFdiffusion-guided framework for the de novo design of catalytic antibody candidates. The framework treats catalytic geometry as a transferable functional unit and uses antibody complementarity-determining regions to reconstruct its surrounding microenvironment. It therefore provides a mechanism-guided and experimentally testable route for introducing enzyme-like chemical functions into the molecular recognition architecture of antibodies.

## Materials and Methods

### Catalytic motif grafting and CDR mapping

The catalytic motif was extracted from ULP1 PROTEASE (PDB: 1EUV), comprising residues His514, Asp531, Gln574, and Cys580 together with the covalently bound ligand l:g. Continuous backbone atoms of the Asp579–Cys580–Gly581 segment served as an alignment anchor. Two acceptor scaffolds were used: the humanized Fv fragment-IGG1-KAPPA 4D5 FAB (PDB: 1FVE, heavy and light chains) and the nanobody-NbBCII10-FGLA (PDB: 3EAK, heavy chain only), both obtained from the RFantibody template library. Complementarity-determining regions (CDRs) were parsed from the HLT (Homology Limiting Templates) remarks in each acceptor PDB.

All combinatorial mappings of the donor catalytic residues onto CDR positions were enumerated, with the constraint that the three-residue anchor fragment must map to a contiguous stretch within a single CDR loop. For each mapping, a Kabsch superposition of the six Cα atoms of the catalytic motif onto the corresponding CDR Cα positions was performed. Candidates were ranked by a composite score incorporating Cα RMSD, local sidechain grafting feasibility, preservation of internal catalytic network distances, and steric clashes with the ligand. The top five candidates for each scaffold were retained. For each selected candidate, the catalytic sidechain terminal atoms and the ligand were placed into the antibody coordinate frame.

### Partial diffusion with RFdiffusion 3

To rebuild the CDR backbone and reconnect the grafted catalytic sidechains, partial diffusion was performed using RFdiffusion 3 [11] with the RFdiffusion 3 _latest.ckpt checkpoint. The input specification defined fixed atoms (catalytic guideposts and the ligand), designable loop regions (CDRs containing the grafted positions), and a partial noise level partial_t = 5.0 (later reduced to 3.0 in motif-scaffolding experiments). Cross-residue bonds were specified to explicitly preserve the covalent linkage between the cysteine sulfur and the ligand carbon C8. Diffusion was run a batch size of 8, generating eight models per candidate.

### Geometric screening of diffused designs

Diffused structures were screened for chain continuity (n_chainbreaks = 0) and for satisfaction of eight catalytic distance constraints. The constraints comprised four ligand-to-residue distances (His-ND1–O1, Asp-OD1–C22, Gln-NE2–O3, Cys-SG–C8) and four inter-residue distances within the catalytic network, all derived from the donor geometry. Distances were evaluated using a custom Python script that parsed mmCIF files through the _atom_site category. Only designs with zero chain breaks and all eight distances within a tolerance of 0.6 Å of their target values progressed to sequence design.

### Sequence design with LigandMPNN

For each geometrically validated diffused structure, non-catalytic CDR residues were redesigned using the LigandMPNN model (ligandmpnn_v_32_010_25.pt) in legacy weights mode. Catalytic residues (His, Asp, Gln, Cys) were excluded from the designable set via a residue whitelist. The ligand was explicitly included as a contextual residue to encourage packing interactions. Sequence generation used a temperature of 0.1, a batch size of 1, and 16 batches per input structure, yielding 16 candidate sequences per diffused model. All designed sequences were verified to preserve catalytic residue identities.

### Structure prediction with RoseTTAFold 3

Designed sequences were folded using RoseTTAFold 3 in template-based mode [12]. The diffused CIF structure served as the template, with both the protein chain(s) and the ligand chain selected for template-guided prediction. Explicit bonds between the catalytic cysteine SG and ligand C8 were supplied to maintain the covalent topology. Five samples per input were generated using 50 denoising steps. Output structures were assessed for global confidence (pLDDT, pTM, ipTM) and for recovery of the eight catalytic distance constraints using the same screening pipeline applied to the diffused models.

### Covalent bond restoration and motif scaffolding

Initial diffusion runs revealed that the cysteine–ligand covalent bond was lost during input preparation, causing ligand displacement in downstream folding. To correct this, the seed PDB files were modified to include explicit CONECT records between the cysteine SG and the ligand C8 atoms, and the RFdiffusion 3 input JSON was updated with a corresponding bonds field. In a subsequent iteration, a stricter motif-scaffolding protocol was applied: partial_t was lowered to 3.0, and additional hotspot constraints were placed on the catalytic sidechain and ligand atoms to enforce their relative geometry during diffusion. LigandMPNN [13] was rerun with the ligand designated as a contextual residue to bias sequence design toward productive ligand-binding pockets. These adjustments successfully preserved the covalent bond in all downstream RoseTTAFold 3 predictions, although non-covalent catalytic contacts remained challenging to fully recapitulate.

## Results

### Catalytic motif grafting and CDR mapping

The donor catalytic motif, comprising residues His514, Asp531, Gln574, and Cys580 together with the covalently bound ligand l:g from the 1EUV crystal structure, was geometrically matched onto the complementarity-determining regions of two antibody scaffolds. The continuous three-residue segment Asp579–Cys580–Gly581 served as the backbone alignment anchor. Exhaustive enumeration yielded 19 632 CDR mappings for the Fv scaffold (hu-4D5-8_Fv.pdb, heavy and light chains) and 306 936 mappings for the nanobody scaffold (h-NbBCII10.pdb, heavy chain only). Candidates were ranked by a composite score incorporating Cα rigid-body superposition RMSD, sidechain grafting feasibility, preservation of internal catalytic network distances, and steric clashes with the ligand, and the top five from each scaffold were retained.

The top-ranked Fv candidate (His514→L3/L91, Asp531→L3/L96, Gln574→H3/H99, Cys580 →H3/H101) exhibited a global Cα RMSD of 3.19 Å and a maximum Cα displacement of 3.95 Å. The top-ranked nanobody candidate (His514 → H3/H107, Asp531→H2/H60, Gln574→H2/H57, Cys580→H3/H105) showed a Cα RMSD of 2.05 Å and a maximum displacement of 2.91 Å, with no coarse ligand clashes, indicating markedly better geometric compatibility.

### RFdiffusion partial diffusion and initial screening

The two best candidates from each scaffold (candidate_001 and candidate_002) were subjected to partial diffusion with RFdiffusion under fixed catalytic atom constraints. A total of 320 independent trajectories were run (80 per candidate) at a partial noise level of partial_t = 5.0, while keeping the terminal atoms of the catalytic sidechains and the entire ligand fixed. All 160 Fv outputs contained one or more chain breaks (100% failure rate), consistent with the larger initial Cα RMSD. For the nanobody candidates, 15 of 80 outputs from candidate_001 (94%) and 16 of 80 outputs from candidate_002 (100%) displayed zero chain breaks, with Cα deviations of only 0.1–0.7 Å, demonstrating that the nanobody framework could accommodate the motif (Figure 1).

**Figure 1.**
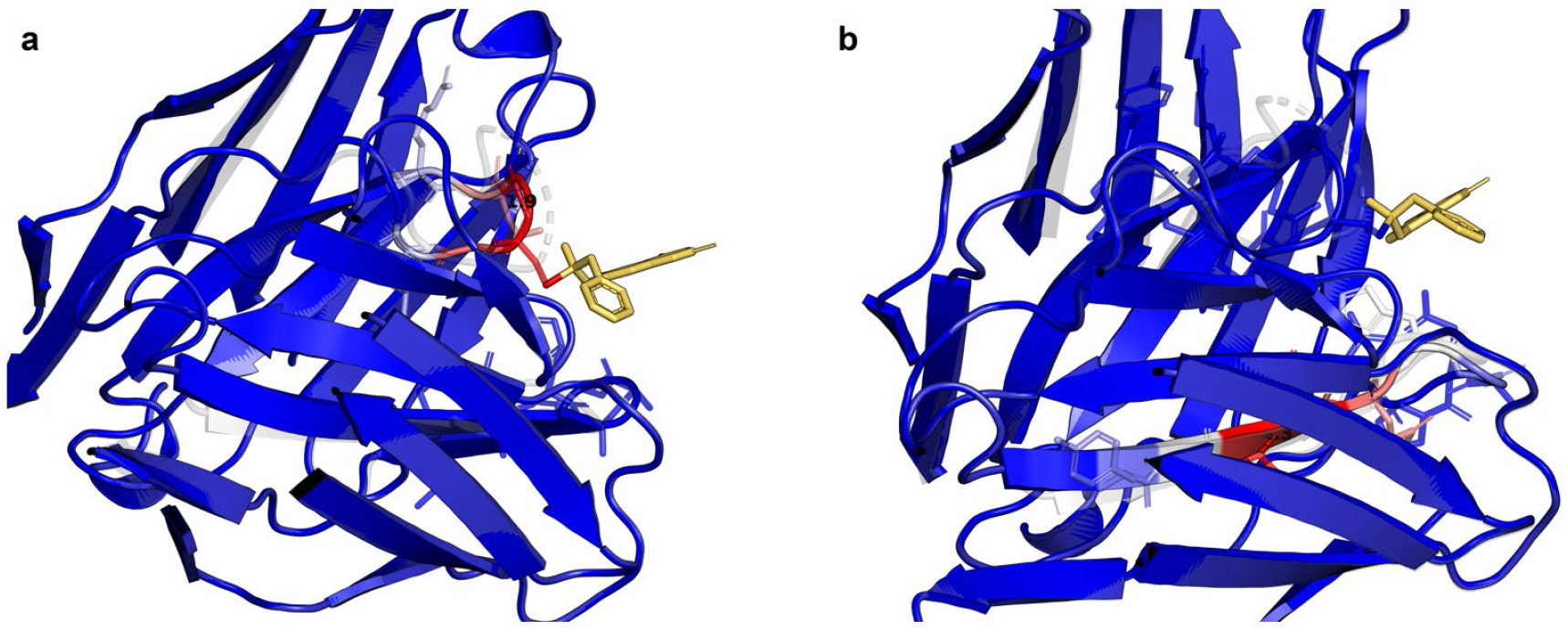
Local backbone discontinuities in RFD3-generated Fv candidates. Seed structures are shown in transparent grey, whereas RFD3 outputs are coloured blue-to-red according to proximity to chain-break hotspots; yellow indicates the fixed ligand. Candidate_001 model_12 shows distortion in CDR-H3 (H100 C–H101 N, 1.913 Å), while candidate_002 model_6 shows an expanded CDR-L3 join point (L88–L89 Cα, 4.295 Å); both models have n_chainbreaks = 2.

After discarding one model with a chain break, 31 chain-break-free structures advanced to catalytic geometry screening. Eight distance constraints were evaluated per candidate: four ligand-to-residue distances (His ND1–O1, Asp OD1–C22, Gln NE2–O3, Cys SG– C8) and four inter-residue distances within the catalytic network (His–Asp, His–Gln, His–Cys, Gln–Cys), with a tolerance of 0.6 Å. Because all catalytic terminal atoms and ligand atoms remained fixed during diffusion, all 31 models satisfied every distance constraint.

For each geometrically validated diffused model, non-catalytic CDR residues were redesigned with LigandMPNN while keeping catalytic residues immutable. Each model produced 16 sequences (temperature 0.1), yielding 496 sequences in total, all of which preserved the identities of the catalytic residues. The top five sequences ranked by ligand-interface sequence recovery (all derived from candidate_001) were selected for all-atom structure prediction with RoseTTAFold 3. Five independent random seeds were run per sequence, producing 25 predicted structures. RoseTTAFold 3 was run in template mode, selecting the nanobody heavy chain and the ligand chain as templates.

The 25 predictions achieved an overall pLDDT of 0.91 and ligand chain pTM values between 0.64 and 0.66. However, catalytic geometry screening revealed that all four ligand-to-residue distances deviated substantially from their targets (measured values reaching 18–30 Å) and the ligand was displaced from the catalytic pocket. Only some inter-residue distances were partially satisfied. No prediction met all eight geometric constraints; the best model passed only two of the eight (Figure 2).

**Figure 2.**
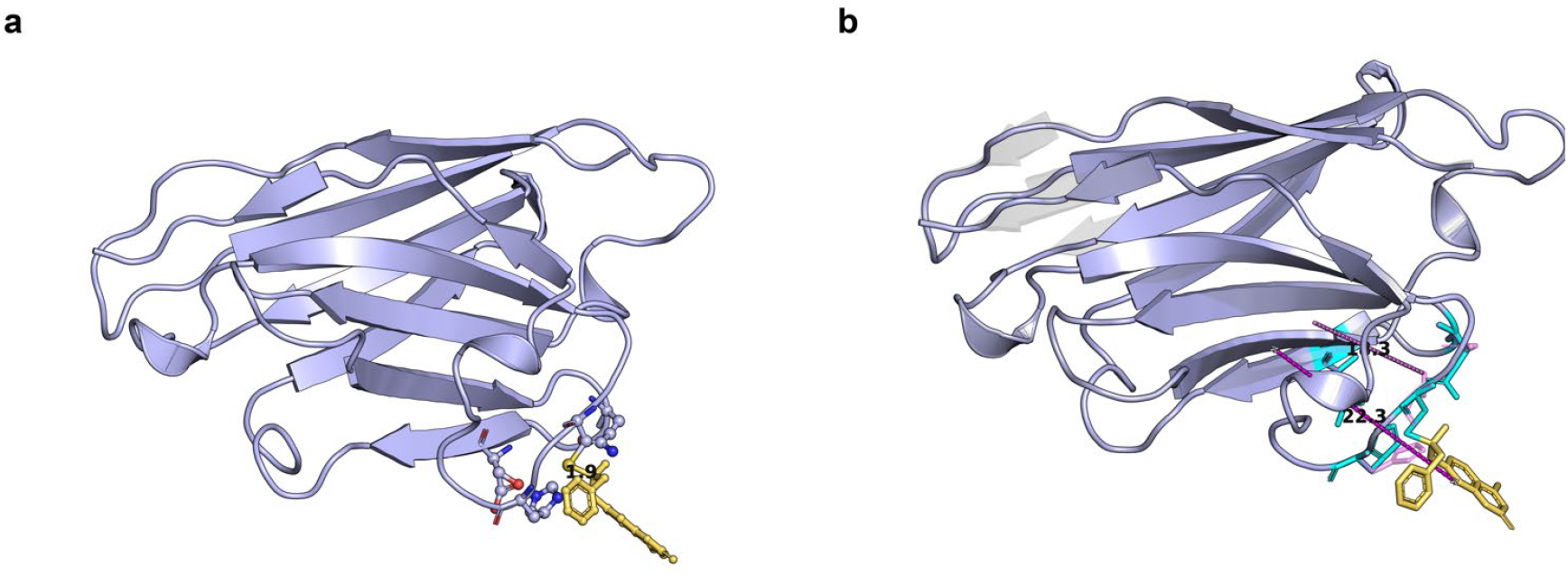
Preserved catalytic guidepost geometry in a nanobody design and ligand displacement during RF3 prediction. (a) RFD3 nanobody candidate_001 model_0 retains a continuous backbone and the fixed His107, Asp60, Gln57 and Cys105 catalytic residues around ligand l:g, with a Cys105 SG– C8 distance of 1.882 Å. (b) After backbone alignment, the unrepaired RF3 prediction displaces the ligand centroid by 22.311 Å and increases the SG–C8 distance to 18.281 Å, despite preserving the overall nanobody fold.

### Diagnosis and repair of the missing covalent bond

Retrospective analysis revealed that the initial seed PDB file, while preserving internal CONECT records for the ligand, lacked the covalent connection between Cys SG and ligand C8. In the donor 1EUV structure this bond measures 1.879 Å and is explicitly encoded by a CONECT record. The missing linkage was not propagated into the RFdiffusion 3 inputs, causing the ligand to be treated as an independent entity during diffusion and subsequently drift in RoseTTAFold 3 predictions due to the absence of covalent topology. To correct this, the following repairs were implemented: (1) explicit CONECT records between Cys SG and ligand C8 were inserted into the seed PDB files; (2) a corresponding bonds field was added to the RFdiffusion 3 input JSON; (3) RFdiffusion 3 partial diffusion was rerun to generate new models with the covalent bond preserved.

The repaired RFdiffusion 3 run produced 16 nanobody models (eight per candidate), all with zero chain breaks. Geometric screening confirmed that the Cys SG–C8 distance recovered to 1.8815 Å for candidate_001 (Δ = 0.002 Å) and 1.8646 Å for candidate_002 (Δ = 0.014 Å), and all eight distance constraints passed.

### Re-evaluation with LigandMPNN and RoseTTAFold 3 after bond repair

The 16 repaired models were again processed through LigandMPNN (16 sequences per model, 256 sequences total), and the top five sequences were subjected to RoseTTAFold 3 prediction (25 structures). The Cys–ligand covalent bond was again explicitly specified in the RoseTTAFold 3 inputs. The covalent Cys SG–C8 distance was maintained in all 25 predictions (1.60–1.95 Å). Nevertheless, the non-covalent catalytic residue-to-ligand distances remained largely unsatisfied; most models still passed only 1–2 of the 8 constraints, indicating that restoring the covalent bond alone was insufficient to reconstruct the full catalytic geometry through template-driven RoseTTAFold 3 folding.

### Enhanced motif constraints and ligand-aware design

To further enforce the local geometry of the catalytic motif, two additional measures were applied. The RFdiffusion 3 partial_t was reduced from 5.0 to 3.0, and a select_hotspots field was introduced to impose extra restraints on the catalytic sidechain and ligand key atoms. In parallel, the LigandMPNN configuration was updated to explicitly include ligand residue information (ligand_residue_numbers) and to enable side-chain context, allowing the sequence design process to be aware of the covalently bound ligand.

A complete RFdiffusion 3–LigandMPNN–RoseTTAFold 3 cycle was rerun under these conditions, yielding 16 diffused models, 256 sequences, and 25 RoseTTAFold 3 predicted structures. The covalent Cys SG–C8 distances remained conserved (1.78– 1.96 Å), but non-covalent catalytic contacts did not improve: across the 25 predictions, the maximum number of satisfied constraints remained at two out of eight, and no structurally complete catalytic geometry was observed.

## Discussion

This study established and systematically evaluated a computational design pipeline for transplanting a catalytic motif from a natural enzyme into the complementarity-determining regions of antibodies. The pipeline encompasses four key stages: geometric matching, partial diffusion, sequence optimization, and all-atom folding validation. The results demonstrated that the nanobody scaffold was significantly superior to the Fv scaffold in geometric compatibility. After RFdiffusion 3 partial diffusion, a series of designs with zero chain breaks and preserved catalytic networks were successfully obtained. However, subsequent RoseTTAFold 3 refolding predictions revealed that although the covalent Cys–ligand linkage could be maintained through explicit topological constraints, the complete non-covalent catalytic contact network could never be recapitulated. This outcome exposes a fundamental limitation of current template-based structure prediction methods in the reconstruction of complex catalytic motifs.

The divergent behaviors of the two scaffolds likely reflect their intrinsic topological constraints. The binding surface of a nanobody resides entirely within a single polypeptide chain. Catalytic residues can be distributed across CDR-H2 and CDR-H3 without requiring coordination across chain interfaces, which substantially reduces the perturbation to the overall scaffold conformation. In contrast, the Fv mapping demands that catalytic residues span both CDR-H3 and CDR-L3 while simultaneously preserving the local continuity of each loop and maintaining the correct relative orientation between the heavy and light chains. This inter-chain dependence increases the difficulty of geometric matching. The larger initial Cα displacements in the two Fv seeds likely severely restricted the conformational freedom available, preventing partial diffusion from repairing the local backbone without introducing chain breaks. Recent diffusion-based antibody design studies have shown that when the generated loops remain compatible with the surrounding framework, the CDRs of both antibodies and nanobodies can support extensive structural remodeling [14]. The present findings extend this principle to a more demanding task, namely accommodating a multi-residue catalytic motif around a small molecule, suggesting that single-chain scaffolds possess an intrinsic advantage in tolerating the structural strain imposed by motif grafting.

The loss of the covalent bond was the core reason for the early failure in this study. The 1.879 Å covalent linkage between Cys SG and ligand C8 in the donor structure was not retained during seed preparation. Consequently, RFdiffusion 3 treated the ligand as an independent entity during diffusion, and RoseTTAFold 3 further mislocalized the ligand due to the absence of topological constraints. By supplementing CONECT records in the seed PDB files and explicitly declaring bonds in the RFdiffusion 3 input JSON, the covalent bond was reliably maintained in all subsequent predictions, with SG–C8 distances consistently falling within the range of 1.60–1.96 Å. This correction confirmed that the correct transmission of covalent topology is indispensable for computational design involving covalent ligands. However, the repair of the covalent bond did not improve the non-covalent catalytic contacts, indicating that the precise shape and chemical complementarity of the ligand-binding pocket cannot be established through topological constraints alone.

To enhance the geometric precision of the catalytic pocket, this study further implemented strategies including lowering the partial diffusion noise level, introducing hotspot constraints, and performing ligand-aware sequence design. Although the sequences generated by LigandMPNN exhibited high ligand-interface sequence recovery and the identities of the catalytic residues remained invariant, the ligand in the RoseTTAFold 3 predictions still deviated from the catalytic site, with deviations of the four catalytic residue–ligand distances commonly exceeding 15 Å. This phenomenon may stem from two factors. First, the template mode of RoseTTAFold 3 relies predominantly on the coordinate bias of the input structure, which itself is a diffusion product. While the ligand position in the input is correct, the surrounding non-covalent interaction network has not been fully optimized. Second, although LigandMPNN can incorporate the ligand as a contextual residue, its training objective is sequence– structure compatibility rather than ligand-binding free energy or catalytic transition-state stabilization. The resulting sequences may therefore not drive folding toward a high-precision catalytic conformation. Furthermore, the non-natural ligand l:g lacks a standard CCD chemical definition, and uncertainties in force-field parameters and protonation states may have further exacerbated the prediction bias.

From the perspective of protein engineering, the fundamental pursuit of catalytic antibodies is not merely to endow antibodies with some enzymatic activity. Rather, it seeks to integrate the programmable molecular recognition ability of antibodies with the catalytic conversion capacity of enzymes into a single protein tool. The traditional strategy is to connect a natural enzyme or a de novo designed enzyme with an antibody in the form of a fusion protein, analogous to the construction of antibody–drug conjugates or immunotoxins [15]. Such “assembled vehicles” achieve the coexistence of targeting and catalytic functions but often suffer from excessive molecular weight, poor structural stability, and a propensity for cleavage at the junction regions [16]. In contrast, the elegance of the catalytic antibody lies in the fact that its catalytic function is directly undertaken by the CDR loops of the antibody itself. The CDR loops serve both as the recognition module and as the catalytic center, eliminating the need for an external targeting module and thereby unifying targeting and catalysis within a single domain.

However, transforming a high-affinity binding pocket into a genuine catalytic pocket requires conditions far more demanding than simple binding. Natural antibodies tend to tightly bind ground-state substrates during evolution. Catalytic function, on the other hand, requires the active center to preferentially stabilize the high-energy transition state while avoiding excessive stabilization of the final reaction product. Otherwise, product inhibition will occur, where the product occupies the active site and cannot be released. A classic solution is to employ transition-state analogs for selection, so that the antibody maintains high affinity for the transition-state analog while its affinity for the reaction product drops rapidly, thereby promoting product turnover. Moreover, the mere presence of catalytic residues does not guarantee catalytic activity. For instance, placing a serine–histidine–aspartate catalytic triad into a CDR only demonstrates the presence of the residue combination. It does not ensure that the pathways for water molecules and proton transfer are rationally arranged. Antibodies naturally achieve broad antigen recognition through the conformational adaptability of their CDR loops, whereas efficient enzymes often require a well-preorganized, rigid active center. If the key catalytic atoms frequently deviate from the catalytic conformation in molecular dynamics simulations, a genuine catalytic capability cannot be realized even if the static structure appears correct.

Based on these considerations, this study introduced a catalytic motif-guided CDR grafting strategy and established a computational design framework for transplanting the key geometric constraints of a natural enzyme active site into a developable antibody scaffold. The framework does not aim to create a fully preorganized rigid active center. Instead, it uses the catalytic geometry of a natural enzyme as a starting point and leverages computational tools to explore the composability of antibody recognition ability and enzyme catalytic function. The framework depends on the accurate preservation of the catalytic motif, the positioning of the substrate (ligand), the optimization of the local microenvironment, and the overall support of the protein scaffold. At present, this study has only focused on the transplantation and validation at the geometric level. It has not yet addressed the evaluation of transition-state stability, proton-transfer pathways, and dynamic behavior. These aspects will define the direction of future research breakthroughs.

Regarding scaffold selection, this study employed a humanized IgG-derived Fv fragment and a nanobody template. It should be noted that immunoglobulin types from different species possess distinct structural features and potential engineering advantages. For example, IgY is the predominant serum antibody type in birds, amphibians, and reptiles, and its antigen-binding fragments exhibit unique structural properties and stability [17]. Designs utilizing humanized antibody scaffolds are more suitable for the direct development of human therapeutic catalytic antibodies. In contrast, designs based on IgY or other non-mammalian antibody scaffolds may show advantages in stability, tissue penetration, and applications targeting specific host environments. The computational grafting methodology established in this work can in principle be extended to these alternative scaffolds, providing flexibility for the development of catalytic antibodies tailored to different species and applications.

From a methodological standpoint, this work achieved the full integration of the pipeline from motif grafting to sequence–structure joint validation and identified the easily overlooked but consequential pitfall of covalent bond loss, offering a troubleshooting pathway for similar studies. At the same time, the results explicitly indicate that relying solely on diffusion models and template-based folding prediction is insufficient for the precise transplantation of catalytic motifs. Potential future improvement directions include introducing a physics-based ligand-binding energy scoring function during the sequence design stage to constrain the sequence space, performing explicit-solvent molecular dynamics simulations on the candidate structures output by RFdiffusion with local relaxation under covalent bond and catalytic distance constraints, or employing transition-state analogs as ligands for de novo pocket generation followed by sequence optimization through Rosetta enzyme design methods. These strategies may help overcome the current bottleneck in reconstructing non-covalent catalytic contacts within the pipeline, ultimately enabling the de novo computational design of catalytic antibodies.

## Data and model availability statement

The dataset and candidate binder models generated in this study are available in the GitHub repository.

## Authors’ contributions

YZ: Conceptualization, Methodology, Investigation, Formal analysis, Visualization, Writing – original draft, Writing – review & editing.

## Reference

1. Tramontano A, Janda KD, Lerner RA: Catalytic antibodies. Science 1986, 234(4783):1566–1570.

2. Pollack SJ, Jacobs JW, Schultz PG: Selective chemical catalysis by an antibody. Science 1986, 234(4783):1570–1573.

3. Lerner RA, Benkovic SJ, Schultz PG: At the crossroads of chemistry and immunology: catalytic antibodies. Science 1991, 252(5006):659–667.

4. Hilvert D: Critical analysis of antibody catalysis. Annual review of biochemistry 2000, 69(1):751–793.

5. Wagner J, Lerner RA, Barbas III CF: Efficient aldolase catalytic antibodies that use the enamine mechanism of natural enzymes. Science 1995, 270(5243):1797–1800.

6. Chiu M, Goulet D, Teplyakov A, Gilliland G: Antibody structure and function: the basis for engineering therapeutics. Antibodies. 2019; 8 (4): 55. In.

7. Paul S, Volle DJ, Beach CM, Johnson DR, Powell MJ, Massey RJ: Catalytic hydrolysis of vasoactive intestinal peptide by human autoantibody. Science 1989, 244(4909):1158–1162.

8. Li X, Liu J, Meng Y, Li J, Zhao J, Liu D, Zhang X: Frontiers in Antibody–Drug Conjugates: Mechanisms, Design Innovations, and Clinical Applications in Targeted Cancer Therapy. Pharmaceuticals 2026, 19(2):324.

9. Chames P, Van Regenmortel M, Weiss E, Baty D: Therapeutic antibodies: successes, limitations and hopes for the future. British journal of pharmacology 2009, 157(2):220–233.

10. Yang W, Wang S, Lee GR, Zhang JZ, Courbet A, Juergens D, Wang X, Schlichthaerle T, Abedi M, Ragotte R et al: The past, present and future of de novo protein design. Nature 2026, 652(8112):1139–1152.

11. Butcher J, Krishna R, Mitra R, Brent RI, Li Y, Corley N, Kim PT, Funk J, Mathis S, Salike S et al: De novo Design of All-atom Biomolecular Interactions with RFdiffusion3. bioRxiv 2025.

12. Corley N, Mathis S, Krishna R, Bauer MS, Thompson TR, Ahern W, Kazman MW, Brent RI, Didi K, Kubaney A et al: Accelerating Biomolecular Modeling with AtomWorks and RF3. bioRxiv 2025.

13. Dauparas J, Lee GR, Pecoraro R, An L, Anishchenko I, Glasscock C, Baker D: Atomic context-conditioned protein sequence design using LigandMPNN. Nat Methods 2025, 22(4):717–723.

14. Luo S, Su Y, Peng X, Wang S, Peng J, Ma J: Antigen-specific antibody design and optimization with diffusion-based generative models for protein structures. Advances in Neural Information Processing Systems 2022, 35:9754–9767.

15. Aboul-Ella H, Gohar A, Ali AA, Ismail LM, Mahmoud AEE-R, Elkhatib WF, Aboul-Ella H: Monoclonal antibodies: from magic bullet to precision weapon. Molecular Biomedicine 2024, 5(1):47.

16. Baldwin E, Schultz PG: Generation of a catalytic antibody by site-directed mutagenesis. Science 1989, 245(4922):1104–1107.

17. Wang H, Zhong Q, Lin J: Egg Yolk Antibody for Passive Immunization: Status, Challenges, and Prospects. J Agric Food Chem 2023, 71(13):5053–5061.

